# Opening Pandora’s Box: Distribution of *Plasmodium* gametocytes in bloodstream

**DOI:** 10.1101/806513

**Authors:** R. Pigeault, J. Isaïa, R. S. Yerbanga, R. D. Kounbobr, J.B. Ouedraogo, A. Cohuet, T. Lefèvre, P. Christe

**Affiliations:** Department of Ecology and Evolution, CH-1015 Lausanne, Switzerland; Institut de Recherche en Sciences de la Santé, Bobo-Dioulasso, Burkina Faso; Unité MIVEGEC, IRD 224-CNRS 5290-Université Montpellier, Montpellier, France

**Keywords:** *P. falciparum*, *P. relictum*, gametocyte density, parasite transmission, diagnosis, *Culex pipiens*

## Abstract

Malaria, a vector borne disease caused by *Plasmodium* spp., remains a major global cause of morbidity and mortality. Optimization of the disease control strategies requires a thorough understanding of the fundamental processes underlying parasite transmission. Although the number of transmissible stages of *Plasmodium* (gametocyte) in human blood is frequently used as an indicator of human-to-mosquito transmission potential, this relationship is not always clear. Important efforts have been made to develop molecular tools to fine-tune gametocyte densities estimation and therefore improve the prediction of mosquito infection rates, but a significant level of uncertainty around this estimate remains. Here we show with both human and avian malaria system that the within-vertebrate host distribution of gametocytes could explain much of this uncertainty. By comparing gametocyte densities in bloodstream between different body parts, we found a difference by nearly 50% in humans and by more than 15% in birds. An estimation of gametocyte density from only one blood sample, as is usually the case, could therefore drastically over- or underestimated the infectivity of gametocyte carriers. This might have important consequences on the epidemiology of the disease since we show, using the avian malaria system, that this variation influences the transmission of the parasite to the mosquito vector. In the light of our results, we argue that it is essential to consider the heterogeneous distribution of gametocyte to improve human diagnosis, identify infectious reservoirs and to test new malaria control strategies.

## Introduction

According to estimates by the World Health Organisation, 219 million cases of malaria had occurred in 2017 with more than 435 000 resulting in death (1). Despite a 50% decline in malaria-related mortality between 2000 and 2015, the number of malaria cases is increasing in several African countries since then. In 2017, the 10 most affected countries on this continent have suffered from 3.5 million additional cases of malaria compared to the previous year. To date, control strategies have aimed at reducing malaria transmission through early human diagnosis and treatment but also through vector control interventions (2). The efficacy of these interventions is however continually challenged and threatened by the evolution of insecticide (3, 4) and drug resistances (5, 6). To overcome resistance issues, the re-emergence of the concept of malaria transmission-blocking strategies (7–10) has boosted the research efforts to find molecules (11, 12) or microorganisms (13–16) able to inhibit the transmission of parasites or to disturb the life cycle of *Plasmodium* in the mosquito vector. Understanding the fundamental processes of parasite transmission from human to mosquito vector is therefore essential to this aim.

A mosquito blood meal represents a volume of 1.5 to 4μl on average (17, 18) and it needs to contain at least one gametocyte (sexual stage) of both sexes to result in malaria infection. Although the shape of the relationship between gametocyte densities measured in the human blood and the probability or the intensity of mosquito infection is not always clear, the likelihood of mosquito infection seems to be mainly dictated by gametocyte density (19, 20). Therefore, the number of gametocytes in human blood is frequently used as an indicator of human-to-mosquito transmission potential (21–23). Robust estimation of gametocyte density is therefore essential to the identification of the human infection reservoir. Although, sensitive molecular techniques have been developed to significantly improve the detection, quantification and possibly sex determination of gametocytes (19, 21, 24–26), the temporal and spatial dynamics of gametocyte distribution and infectivity in vertebrate hosts remain relatively neglected. Gametocyte density are mostly estimated from a single blood sample from a single body location (e.g. finger prick or antecubital venous blood). Such snapshots certainly fail to grasp the complex and dynamic nature of vertebrate host infection. For instance, a recent study showed that at night, rodent malaria *P. chabaudi* gametocytes are twice as infective compared to the daily ones, despite being less numerous in the blood (27). In the avian malaria system, a periodic increase of parasitaemia is observed in the late afternoon (28). Regarding the spatial distribution of mature gametocytes in the bloodstream, a handful of studies have compared their density between venous and capillary blood. A majority of them have found a higher density of gametocytes in capillary than in venous blood ((29–33) but see (34, 35)). However, comparisons of the number of gametocytes from the same blood compartment (veins or capillaries) but from different body parts has been investigated in a unique study conducted in 1952, where a 3-fold higher prevalence of gametocytes in skin capillary blood compared to thick smears prepared from finger-prick has been observed (36).

Mature gametocytes have long been considered passively displaced by the blood flow (37–39), suggesting a random or even a homogeneous distribution in the peripheral circulatory compartment (40–44). A study carried out in the early 2000s has shaken this paradigm by providing evidences that the distribution of gametocytes ingested by mosquitoes from the skin of three naturally-infected volunteers followed a negative binomial distribution and not a Poisson distribution, as expected if gametocytes circulate in a homogenous pattern in the peripheral bloodstream (45). This result suggests an aggregated distribution of gametocytes in the vertebrate peripheral bloodstream (45, 46). Another indirect evidence of the non-homogeneous distribution of gametocytes in the vertebrate host body comes from a review of the literature. In human malaria, the proportion of studies showing a positive relationship between gametocyte density and parasite transmission to mosquito increases drastically when mosquitoes were fed via an artificial membrane-feeding method compared to mosquitoes fed directly through the skin of an infected host (Figure S1, Table S1). Artificial feeding techniques eliminates the potential heterogeneity in gametocyte spatial distribution in the host and the density of gametocytes is measured from the blood that will then be used to feed mosquitoes.

To date, no study has empirically studied the distribution of *Plasmodium* gametocytes in the peripheral blood compartment of the vertebrate host. Our specific aims in this study are to answer two questions. First, is the *Plasmodium* gametocytes density homogeneous or heterogeneous between different body parts of the vertebrate host? Second, is the transmission of *Plasmodium* to the vector comparable according to the location of mosquito bites? This work uses both human (*P. falciparum/Anopheles gambiae s.s*) and avian malaria (*P. relictum/Culex pipiens*) systems to measure simultaneously gametocyte density at two different locations of host body: the left and right hand in humans and the left and right leg in birds. The intra-individual variation rate in gametocyte density was used as a proxy to determine whether *Plasmodium* sexual stage is homogeneously distributed in host body. Due to ethical reasons, the impact of mosquito biting sites on parasite transmission was carried out only with the avian malaria system. Avian malaria is the oldest experimental model for investigating the life cycle of *Plasmodium* parasites and it was rapidly identified as the ideal experimental system for understanding the biology of human malaria parasites (47). Hereafter, we show that both *P. falciparum* and *P. relictum* are not homogeneously distributed in the peripheral bloodstream of the host and we argue that this can have implications for the parasite transmission to the mosquito vector.

## Results

### Spatial heterogeneity of *Plasmodium* infection in vertebrate host

The number of gametocytes was significantly different between the two hands 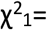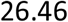, P < 0.0001, Figure 1A). Nevertheless, it was not always the same hand (right or left) that had the highest gametocyte density (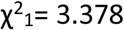, P = 0.077). The average variation rate of the number of gametocytes between the two parts of the body was 0.45 ± 0.06 (Figure 1A) and variation rates were negatively correlated to human gametocytaemias (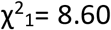, P = 0.003, Figure 2A). Fitting the quadratic term (gametocytaemia^2^) improved the model fit (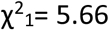, P = 0.017), suggesting that variation rate was a decelerating polynomial function of human gametocytaemia (Figure 2A).

**Figure 1.**
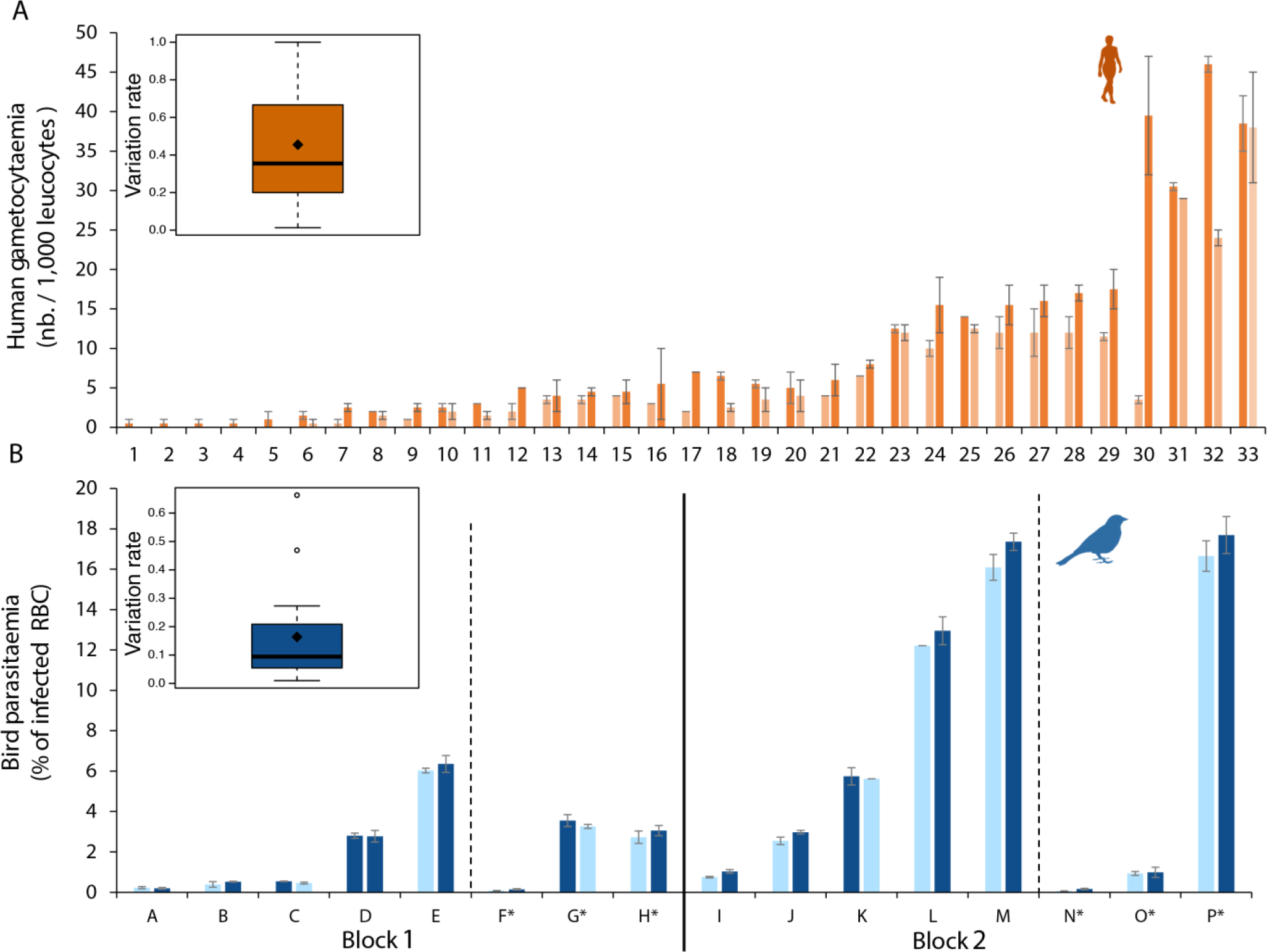
Variation in infection density between two body parts. (A) Variation in human gametocyte density between the left hand (left bar) and the right hand (right bar). (B) Variation in bird parasite density between the left leg (left bar) and the right leg (right bar). The black vertical line on panel B separates the two experimental blocks (see materials & methods). Each number (Human) or letter (bird) correspond to one individual. * corresponds to control individuals (unexposed to mosquito bites). Light color bar correspond to the body part with the lower gametocyte density (human) or parasite density (bird), the dark color bar correspond to the body part with the higher gametocyte (human) or parasite (bird) density. Boxplots represent the variation rate in parasite density measured between the two body parts. Error bars represent standard error around the mean. RBC: red blood cell.

**Figure 2.**
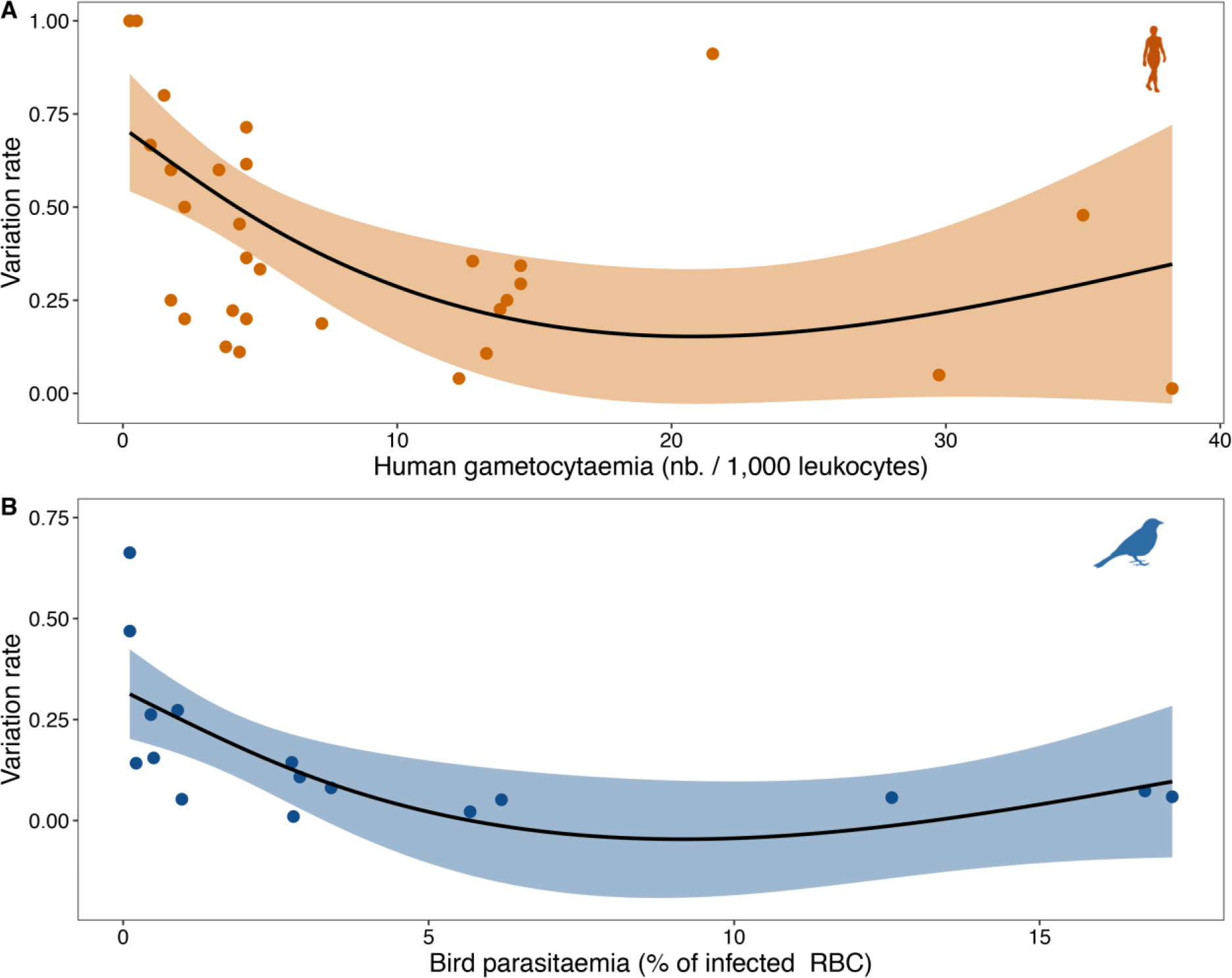
Relationship between variation rate and average parasite density. (A) Variation rate of gametocyte density between the left and the right hand in human. (B) Variation rate of parasite density between the left and the right leg in bird. Shaded areas on either side of the regression line represent the 95% CI.

Bird parasitaemia was used as a proxy for gametocytemia because parasitaemia and gametocytaemia are strongly positively correlated in this system (see Figure 2 in Pigeault et al. 2015). Bird parasitaemia was significantly different between the two legs (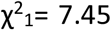, P = 0.006, Figure 1B), but it was not always the same body part (right or left leg) that showed the highest parasite density (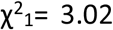, P = 0.082). Since some of the birds were exposed to mosquitoes before blood sampling (see below), we also compared the parasitaemia of control and exposed birds and we did not observe any difference (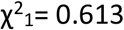, P = 0.433). The average variation rate of parasitaemia between the two legs was 0.16 ± 0.04 (Figure 1B). As for human malaria, the variation rates were negatively correlated to bird parasitaemias (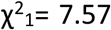, P = 0.006, Figure 2B) and fitting the quadratic term (parasitaemia^2^) improved the model fit (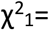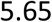, P = 0.017, Figure 2B).

### Parasite transmission to mosquito vector

To investigate the effect of the location of mosquito bites on *Plasmodium* transmission, left and right legs of infected birds (*Serinus canaria*) were independently but simultaneously exposed to mosquitoes (*Culex pipiens*) during 3 hours. Immediately after the exposure session, blood from both legs was collected to measure parasite densities and each leg was classified as either lower infected leg (LIL) or higher infected leg (HIL). Blood-fed mosquitoes were dissected one-week post-blood meal to count the number of parasites in their midgut.

There was no significant difference between female mosquitoes fed on the lower (LIL) or on the higher infected leg (HIL) in either the proportion of blood-fed females (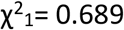 P = 0.406), blood meal size (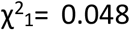 P = 0.826) and infection prevalence (proportion of mosquitoes containing at least 1 oocyst, 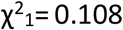 P = 0.478, Figure 3A). The analysis of oocyst burden included mosquitoes having at least one oocyst in the midgut. A significant difference in oocyst burden was observed (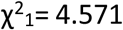 P = 0.032, Figure 3B). Females fed on the HIL had a higher oocyst burden than females fed on the LLP (mean ± s.e. females fed on LLP: 9.75 ± 2.47, females fed on LHP: 16.27 ± 2.65).

**Figure 3.**
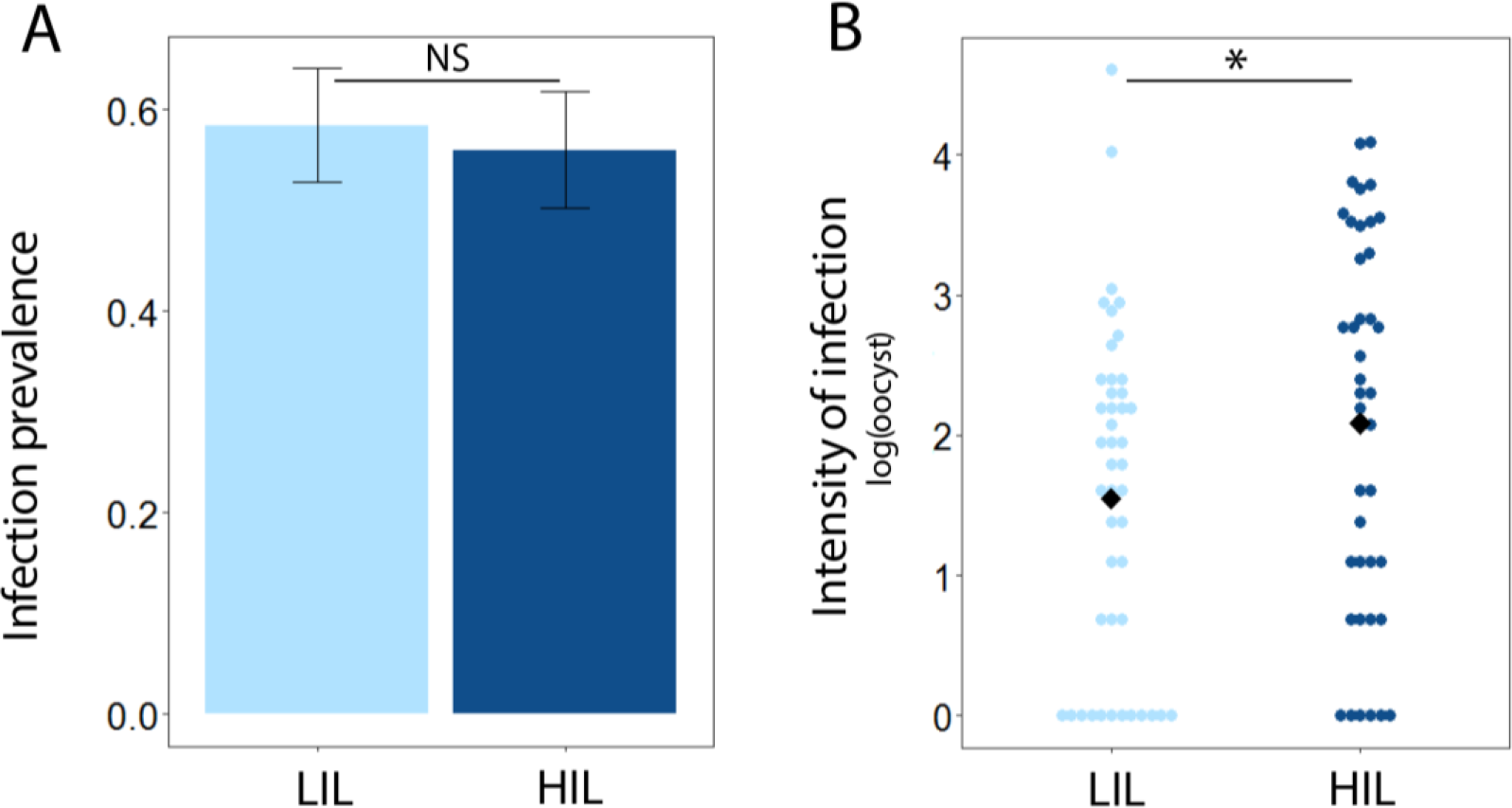
Mean infection prevalence (A) and oocyst burden (B) in mosquitoes fed on either the lower infected leg (LIL) or on the higher infected leg (HIL). Bars represent standard error around the mean.

## Discussion

Reduction of *Plasmodium* transmission from the vertebrate host to the insect vector is a key component of global efforts to control malaria (48). Understanding the processes that underlie the relationship between *Plasmodium* gametocyte densities and mosquito infection is therefore crucial to assess the effectiveness of control programs and their effects on transmission. Nevertheless, an essential first step to meet this goal is to obtain an accurate estimation of gametocyte density in infected host. For this purpose, significant efforts have been made to develop new molecular tools to detect and estimate more precisely the gametocyte densities for both males and females. It is interesting to note that these methods have also significantly improved the prediction of mosquito infection rates (19). Nevertheless, a significant level of uncertainty remains.

An aspect that is all too rarely mentioned and, above all, very rarely quantified concerns the spatial distribution of mature gametocytes within the vertebrate host. Here, we showed that the number of gametocytes fluctuates by nearly 50% between the two human hands and by more than 15% between the two legs of the birds. Therefore, gametocytes did not seem to be homogeneously distributed within the vertebrate host body and we highlight that this could have an impact on parasite transmission to the mosquito vector. Using the avian malaria system, we showed that mosquitoes fed on the least infected body part have a lower parasite burden than those fed on the most infected part. Consequently, using a single measurement of gametocyte density from a single blood sample at a unique body part does not provide a good estimate of a host’s infectivity. Our results suggest that the most efficient way to obtain a more accurate estimate of the total gametocyte densities and therefore a more predictable infectiousness indicator would be to combine several independent density measurements from different body parts.

The mechanisms leading to the establishment of a non-homogeneous distribution of *Plasmodium* in the blood of the vertebrate host are unknown. Gametocytes are not motile and cannot therefore actively migrate to accumulate in the capillaries. Passive accumulation of gametocytes in some sub-dermal capillaries could induce a non-homogeneous distribution of *Plasmodium* in the vertebrate host. For instance, the elongated asymmetric curvature of *P. falciparum* gametocytes may facilitate their blockage in the dermal capillaries (49). Mature gametocyte aggregation might also partly explain the spatial heterogeneity in the distribution of gametocytes (45, 50). Active aggregation mechanisms with, for instance binding interaction between infected red blood cells containing late developmental stages of the gametocyte, have however never been observed in both human and avian malaria parasite (“rosetting-like” adhesion, (51)).

When the gametocyte density is low, an adaptive strategy allowing the aggregation of several sexual stages of *Plasmodium* may increase the probability that a mosquito will be infected (52). In this case, while the majority of blood-fed mosquitoes did not ingest any parasite, those biting an area containing aggregated gametocytes will be undoubtedly infected by malaria. Nonetheless, this strategy would be non-adaptive if the gametocyte density increases to a level that allows the ingestion of at least two gametocytes regardless of the mosquito biting site. In this case, homogeneous distribution should optimize transmission. A plastic strategy with a regulation of the level of aggregation according to vertebrate host gametocytaemia would then be an optimal process to promote the transmission of the parasite to the mosquito vector throughout the infection. Interestingly, our results fit in with a plastic adaptive strategy. We showed that the variation rate in the number of gametocytes count between two different body parts decreases with the increase in average gametocytaemia.

Despite the match between our empirical results and the plastic adaptive strategy hypothesis, alternative explanations challenge our results. Given that malaria infection is temporally dynamic (28, 53), the single measurement used in this study to compare the number of parasites between different body parts does not allow to determine whether non-homogeneous distribution is stable over time or whether it is a single snapshot that does not reflect a more complex dynamic process. It would be relevant to monitor gametocyte density at different body parts with repeated measurements over the course of the infection. Of particular relevance would be to compare gametocytes densities among different body locations in regard to variation in mosquito attraction to these sites. For instance, it is known that the major vectors of *P. falciparum* (*An. gambiae s.s.*, *An. arabiensis*, *An. funestus*) all have strong preference for feeding close to the ground which is associated to increased biting rate on legs, ankles and feet (54–56). Accordingly, higher gametocyte density would be expected in these highly attractive body parts.

Improving the detection and estimation of gametocyte density in infected hosts is fundamental to improve diagnosis of gametocyte carriers and therefore identify infectious reservoirs but also to develop and test new malaria control strategies. In this study, we found that the gametocyte burden varies significantly between different body parts. We argue that it would then be essential to perform several blood samples at different body parts with respect to preference for biting to refine our understanding of the within-host malaria infection dynamic and ultimately the fundamental processes underlying the parasite transmission from human-to-mosquito.

## Materials & Methods

### Human malaria

The study was conducted at the Institut de Recherche en Sciences de la Santé in Bobo Dioulasso, South-Western Burkina Faso. The intensity of malaria transmission is high and perennial in this area with a peak from August to November. Blood slides were collected from December 2018 to July 2019 from asymptomatic children aged 5-12 years attending the elementary schools of Dandé, Soumousso, Klesso, Samandeni - four villages located in the surroundings of Bobo Dioulasso. *P. falciparum* is the predominant parasite species in these villages, accounting for more than 95% of malaria cases (57).

Samples were collected prior to treatment with a dose of artemether–lumefantrine according to National Malaria Control Programme recommendation and after written informed consent was obtained from the parent(s) or guardian(s). Ethical clearance was provided by the national ethics committee of Burkina Faso (no. 2018-9-118) and the institutional committee of IRSS (no. A06-2018/CEIRES).

Finger prick blood samplings were carried out on the two hands of each volunteer so that two Giemsa-stained blood smears per volunteer were screened for asexual parasites and gametocytes. Gametocyte density was estimated by microscopy against 1000 leucocytes and slides were declared negative after a minimum reading of 100 fields. Each slide was read twice by two independent qualified microscopists (57). The two gametocyte density measurements for each slide were then used to calculate an average gametocyte density for each hand of each individual. The two hands (right and left) of each individual were then classified as lower infected hand or higher infected hand.

### Avian malaria

This study was approved by the Ethical Committee of the Vaud Canton veterinary authority, authorization number 1730.4.

#### Parasite strain

*Plasmodium relictum* is the most prevalent form of avian malaria in Europe (58). The lineage used in these experiments (lineage SGS1) was isolated from infected great tits (*Parus major*) caught in the region of Lausanne (Switzerland) in 2015. Parasite was passaged to an uninfected canary (*Serinus canaria*) by intraperitoneal injection and has been maintained by carrying out regular passages between infected and naïve canaries. For both experimental blocks, eight uninfected canaries were experimentally inoculated by means of an intraperitoneal injection of 150_200μL of an infected blood pool constituted of a mixture of blood from five infected canaries. Birds of the same experimental block were infected with the same pool of blood. The eight infected birds of each block were assigned to two treatments: “exposed” (n=5) or “unexposed” (n=3) to mosquito bites (Figure 1B).

#### Mosquitoes rearing

The two experimental blocks were conducted with wild *Culex pipiens* mosquito collected from the field (Lausanne, 46°31’25.607”N 6°34’40.714”E, altitude: 380 m) and maintained under laboratory condition since August 2017. Mosquitoes were reared as described by Vézilier et al. (59) in an insectary at 25°C ± 1°C, 70 ± 5% RH and with 14L:10D photoperiod. One day before each experimental block, 500 7-10 days old female mosquitoes were haphazardly chosen from different emergence cages and placed inside new cages (100 females per cage). Females were deprived of sugar solution for 24h to increase hunger levels in order to maximize overall biting rate. Water was provided from 24h to 6h before the experiment to prevent dehydration.

#### Experimental design

The two experimental blocks were carried in February and April 2018 respectively. Twelve days post-bird infection (to coincide with the acute phase of the *Plasmodium relictum* infection in canaries, (60)) “exposed” individuals were placed individually in two compartments cages designed for physically separating their two legs. At 6:00 pm, a batch of 45-50 uninfected female mosquitoes was added in each compartment (left and right) for 180 minutes. Unexposed birds were placed in the same experimental condition but without mosquitoes. At the end of the mosquito exposure session (9:00 pm) a red lamp was used to capture mosquitoes and five microliters of blood were collected from the medial metatarsal vein of both canary legs. For each leg of each bird, three drops of blood were smeared onto three different microscope slides. Blood fed mosquitoes were placed individually into numbered plastic tube covered with a mesh. Food was provided in the form of a cotton pad soaked in a 10% sugar solution placed on top of each tube. Between 7 and 8 days post-blood meal, female mosquitoes were dissected and the number of *Plasmodium* oocysts in their midgut was counted with the aid of a binocular microscope (59). Haematin excreted at the bottom of each plastic tube was quantified as an estimate of the female’s blood meal size (59).

The intensity of bird infection (parasitaemia) was determined visually by calculating the number of infected red blood cell per 3000 erythrocytes in randomly chosen fields on the blood smears (58). Parasitaemia measured on the three blood smears were used to calculate an average parasitaemia (mean ± SE) for each leg of each bird. The two legs (right and left) of each bird was then classified as Lower infected leg (LIL) or higher infected leg (HIL). All parasitaemia was measured by the same experimenter. Parasitaemia was used as a proxy for transmissible stage (gametocytes) production because parasitaemia and gametocytaemia are strongly positively correlated in this system (see Figure 2 in (60)).

### Statistical analyses

#### Human malaria

The analyses of the number of gametocytes were carried out using a generalized linear mixed model (GLMM) procedure with a negative binomial family. To study whether gametocyte numbers were similar between both human hands, the average gametocyte count was used as response variable. Hand class (lower or higher infected) and hand side (left or right) were fitted as fixed factor and individual were fitted as random factor.

#### Avian malaria

The analyses of birds’ parasitaemia were carried out using a linear mixed model procedure with a normal error distribution. To study whether parasitaemia was similar between both legs, the average parasitaemia was used as response variable. Leg class (lower or higher infected), leg side (left or right) and bird treatment (exposed or unexposed to mosquito) were fitted as fixed factors. Individual, nested in experimental block, was fitted as random factor.

Mixed effects models were used to analyze the effect of bird leg (LLP or LHP) on the mosquito blood meal rate (proportion of females that had taken a blood meal), blood meal size and *Plasmodium* transmission to the vector. Explanatory variables, leg class (LLP or LHP) and haematin (when it was appropriate), were fitted as fixed factors. Individuals, nested in experimental block, were fitted as a random factor. Blood meal rate and infection prevalence (oocyst presence/absence) were analyzed using GLMM with a binomial error distribution (lme4 package). Blood meal size and mosquito infection intensity (number of oocysts) were analyzed using lmer with normal error distribution (lme4 package). For the analysis of infection intensity, only individuals that developed ≥ 1 oocyst were included.

Maximal models, including all higher order interactions, were simplified by eliminating non-significant terms and interactions to establish a minimal model (61). Non-significant interactions and terms were removed step by step according to their significance using a likelihood ratio test which is approximately distributed as a Chi square distribution (0.05 was used as the cutoff for p-value significance, (62)). The significant Chi square-values given in the text are for the minimal model, while non-significant values correspond to those obtained before deletion of the variable from the model. Analyses were carried out using the R statistical package (v. 3.4.1, http://www.cran.r-project.org/).

---

**Figure S1.**
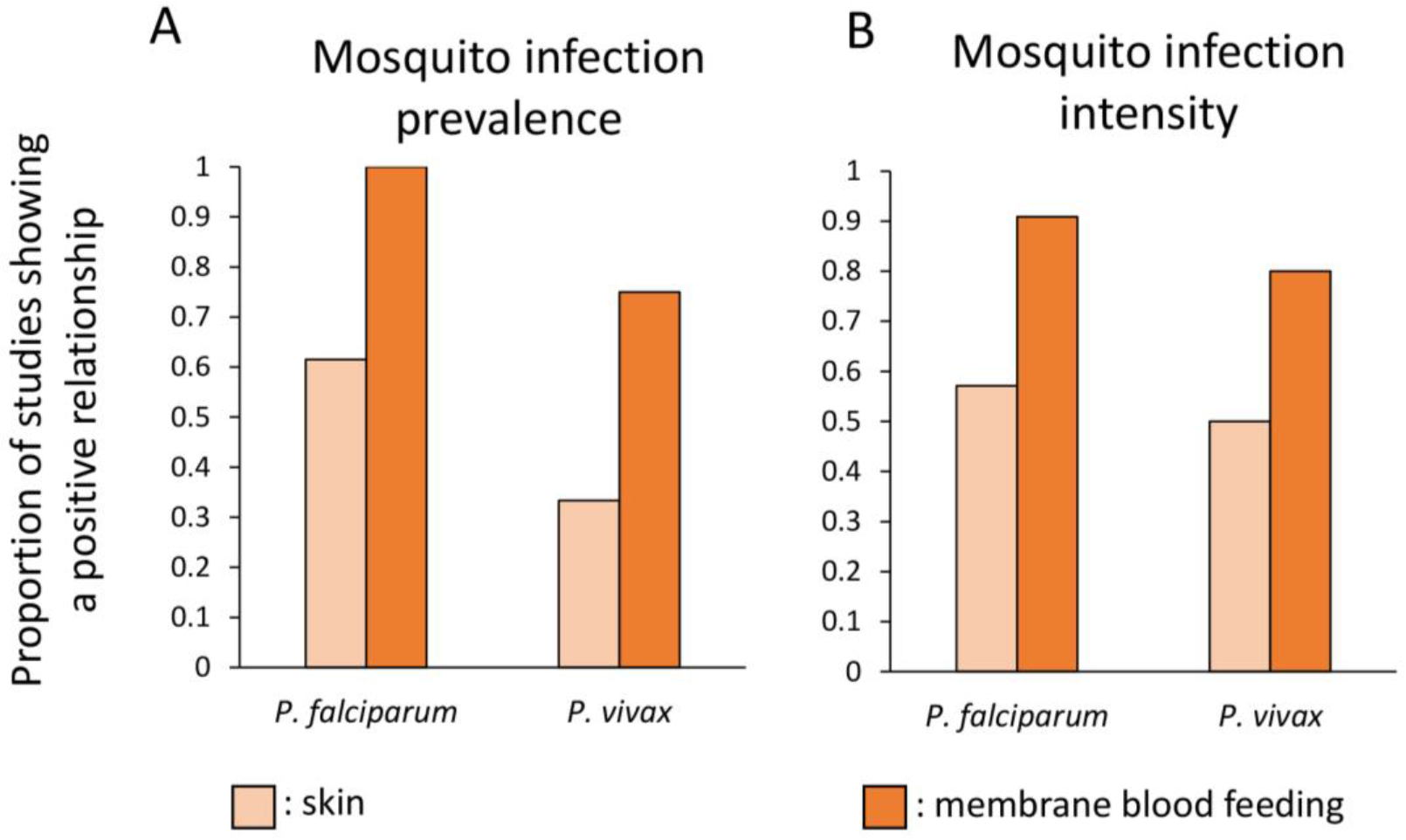
Proportion of studies showing a positive relationship between gametocyte density, estimated from human blood, and (A) mosquito infection prevalence or (B) oocyst burden. The light-orange bars represents mosquitoes fed directly on the skin of infected individuals, the dark-orange bars represents mosquitoes fed with artificial membrane feeding. See Table S1 for references.

## References

1. WHO | World malaria report 2018. WHO (September 27, 2019).

2. S. Bhatt, et al., The effect of malaria control on *Plasmodium falciparum* in Africa between 2000 and 2015. Nature 526, 207–211 (2015).

3. M. Weill, et al., Comparative genomics: Insecticide resistance in mosquito vectors. Nature 423, 136–137 (2003).

4. H. Ranson, N. Lissenden, insecticide resistance in african *Anopheles* mosquitoes: A worsening situation that needs urgent action to maintain malaria control. Trends Parasitol. 32, 187–196 (2016).

5. R.N. Price, et al., Mefloquine resistance in *Plasmodium falciparum* and increased pfmdr1 gene copy number. Lancet Lond. Engl. 364, 438–447 (2004).

6. F. Ariey, et al., A molecular marker of artemisinin-resistant *Plasmodium falciparum* malaria. Nature 505, 50–55 (2014).

7. R. Carter, D.H. Chen, Malaria transmission blocked by immunisation with gametes of the malaria parasite. Nature 263, 57 (1976).

8. R. Carter, Transmission blocking malaria vaccines. Vaccine 19, 2309–2314 (2001).

9. O.K. Doumbo, K. Niaré, S.A. Healy, I. S. and P.E. Duffy, Malaria transmission-blocking vaccines: present status and future perspectives. Malar. Elimin. - Leap Forw. (2018) https://doi.org/10.5772/intechopen.77241 (September 6, 2019).

10. F.K. Acquah, J. Adjah, K.C. Williamson, L.E. Amoah, Transmission-blocking vaccines: old friends and new prospects. Infect. Immun. 87, e00775–18 (2019).

11. D.K. Mathias, et al., A small molecule glycosaminoglycan mimetic blocks *Plasmodium* invasion of the mosquito midgut. PLOS Pathog. 9, e1003757 (2013).

12. G. Niu, et al., Targeting mosquito FREP1 with a fungal metabolite blocks malaria transmission. Sci. Rep. 5, 14694 (2015).

13. Y. Dong, F. Manfredini, G. Dimopoulos, Implication of the mosquito midgut microbiota in the defense against malaria parasites. PLoS Pathog 5, e1000423 (2009).

14. C.M. Cirimotich, J.L. Ramirez, G. Dimopoulos, Native microbiota shape insect vector competence for human pathogens. Cell Host Microbe 10, 307–310 (2011).

15. G.L. Hughes, A. Rivero, J.L. Rasgon, *Wolbachia* can Enhance *Plasmodium* infection in mosquitoes: implications for malaria control? PLOS Pathog. 10, e1004182 (2014).

16. A. Boissière, et al., Midgut microbiota of the malaria mosquito vector *Anopheles gambiae* and interactions with *Plasmodium falciparum* infection. PLoS Pathog 8, e1002742 (2012).

17. M.J. Klowden, A.O. Lea, Blood meal size as a factor affecting continued host-seeking by *Aedes aegypti* (L.). Am. J. Trop. Med. Hyg. 27, 827–831 (1978).

18. A. Ogunrinade, The measurement of blood meal size in *Aedes aegypti* (L.). Afr. J. Med. Med. Sci. 9, 69–71 (1980).

19. J. Bradley, et al., Predicting the likelihood and intensity of mosquito infection from sex specific *Plasmodium falciparum* gametocyte density. eLife 7(2018).

20. D.F. Da, et al., Experimental study of the relationship between *Plasmodium* gametocyte density and infection success in mosquitoes; implications for the evaluation of malaria transmission-reducing interventions. Exp. Parasitol. 149, 74–83 (2015).

21. A. Ouédraogo, et al., Dynamics of the human infectious reservoir for malaria determined by mosquito feeding assays and ultrasensitive malaria diagnosis in Burkina Faso. J. Infect. Dis. 213, 90–99 (2016).

22. C. Koepfli, G. Yan, *Plasmodium* gametocytes in field studies: Do we measure commitment to transmission or detectability? Trends Parasitol. 34, 378–387 (2018).

23. H.C. Slater, et al., The temporal dynamics and infectiousness of subpatent *Plasmodium falciparum* infections in relation to parasite density. Nat. Commun. 10, 1433 (2019).

24. E. Essuman, et al., A Novel gametocyte biomarker for superior molecular detection of the *Plasmodium falciparum* infectious reservoirs. J. Infect. Dis. 216, 1264–1272 (2017).

25. M. Gruenberg, et al., Molecular and immunofluorescence-based quantification of male and female gametocytes in low-density *P. falciparum* infections and their relevance for transmission. J. Infect. Dis. (2019) https://doi.org/10.1093/infdis/jiz420.

26. L. Meerstein-Kessel, et al., A multiplex assay for the sensitive detection and quantification of male and female *Plasmodium falciparum* gametocytes. Malar. J. 17, 441 (2018).

27. P. Schneider, et al., Adaptive plasticity in the gametocyte conversion rate of malaria parasites. PLOS Pathog. 14, e1007371 (2018).

28. R. Pigeault, Q. Caudron, A. Nicot, A. Rivero, S. Gandon, Timing malaria transmission with mosquito fluctuations. Evol. Lett. (2018) https://doi.org/10.1002/evl3.61.

29. R.E. Sinden, “Sexual Development of Malarial Parasites” in Advances in Parasitology, J.R. Baker, R. Muller, Eds. (Academic Press, 1983), pp. 153–216.

30. J. Ouedraogo, T. Guiguemde, A. Gbary, Etude comparative de la densite parasitaire de Plasmodium falciparum dans le sang capillaire et dans le sang veineux chez des porteurs asymptomatiques (région de Bobo-Dioulasso, Burkina Faso). Médecine D’Afrique Noire 38, 601–605 (1991).

31. A. Njunda, N. Assob, S. Nsagha, M. Mokenyu, E. Kwenti, Comparison of capillary and venous blood using blood film microscopy in the detection of malaria parasites: A hospital based study. Sci J Microbiol 2, 89–94.

32. K. Kast, et al., Evaluation of *Plasmodium falciparum* gametocyte detection in different patient material. Malar. J. 12, 438 (2013).

33. J. Mischlinger, et al., Use of capillary blood samples leads to higher parasitemia estimates and higher diagnostic sensitivity of microscopic and molecular diagnostics of malaria than venous blood samples. J. Infect. Dis. 218, 1296–1305 (2018).

34. M.M. Sandeu, et al., Do the venous blood samples replicate malaria parasite densities found in capillary blood? A field study performed in naturally-infected asymptomatic children in Cameroon. Malar. J. 16, 345 (2017).

35. A. Lehane, et al., Comparison on simultaneous capillary and venous parasite density and genotyping results from children and adults with uncomplicated malaria: a prospective observational study in Uganda. BMC Infect. Dis. 19, 559 (2019).

36. L. van den Berghe, M. Chardome, E. Peel, Supériorité des préparations de scarification du derme sur les préparations de sang périphérique pour le diagnostic de malaria. Instit Med Trop 9, 553–562 (1952).

37. C.G. Huff, W. Bloom, A malarial parasite infecting all blood and blood-forming cells of birds. J. Infect. Dis. 57, 315–336 (1935).

38. C.J. Janse, et al., In vitro formation of ookinetes and functional maturity of *Plasmodium berghei* gametocytes. Parasitology 91, 19–29 (1985).

39. R.G. Douglas, R. Amino, P. Sinnis, F. Frischknecht, Active migration and passive transport of malaria parasites. Trends Parasitol. 31, 357–362 (2015).

40. M.F. Boyd, W.K. Stratman-Thomas, S.F. Kitchen, On the relative susceptibility of *Anopheles quadrimaculatus* to *Plasmodium vivax* and *Plasmodium falciparum*. Am. J. Trop. Med. Hyg. s1-15, 485–493 (1935).

41. R.C. Muirhead-Thomson, E.C. Mercier, Factors in malaria transmission by *Anopheles albimanus* in Jamaica. Ann. Trop. Med. Parasitol. 46, 103–116 (1952).

42. L.H. Miller, H.N. Fremount, S.A. Luse, Deep vascular schizogony of *Plasmodium knowlesi* in *Macaca mulatta*. Am. J. Trop. Med. Hyg. 20, 816–824 (1971).

43. M.E. Smalley, S. Abdalla, J. Brown, The distribution of *Plasmodium falciparum* in the peripheral blood and bone marrow of Gambian children. Trans. R. Soc. Trop. Med. Hyg. 75, 103–105 (1981).

44. M. Pritsch, et al., Stability of gametocyte-specific Pfs25-mRNA in dried blood spots on filter paper subjected to different storage conditions. Malar. J. 11, 138 (2012).

45. G. Pichon, H.P. Awono-Ambene, V. Robert, High heterogeneity in the number of *Plasmodium falciparum* gametocytes in the bloodmeal of mosquitoes fed on the same host. Parasitology 121 (Pt 2), 115–120 (2000).

46. F.O. Gaillard, C. Boudin, N.P. Chau, V. Robert, G. Pichon, Togetherness among *Plasmodium falciparum* gametocytes: interpretation through simulation and consequences for malaria transmission. Parasitology 127, 427–435 (2003).

47. A. Rivero, S. Gandon, Evolutionary ecology of avian malaria: Past to present. Trends Parasitol. (2018) https://doi.org/10.1016/j.pt.2018.06.002 (June 26, 2018).

48. P.L. Alonso, et al., A Research agenda to underpin malaria eradication. PLOS Med. 8, e1000406 (2011).

49. M. Nacher, Does the shape of *Plasmodium falciparum* gametocytes have a function? Med. Hypotheses 62, 618–619 (2004).

50. C.P. Nixon, *Plasmodium falciparum* gametocyte transit through the cutaneous microvasculature: A new target for malaria transmission blocking vaccines? Hum. Vaccines Immunother. 12, 3189–3195 (2016).

51. G. Neveu, et al., *Plasmodium falciparum* gametocyte-infected erythrocytes do not adhere to human primary erythroblasts. Sci. Rep. 8, 1–11 (2018).

52. R. E. L. Paul, S. Bonnet, C. Boudin, T. Tchuinkam, V. Robert, Aggregation in malaria parasites places limits on mosquito infection rates. Infect. Genet. Evol. 7, 577–586 (2007).

53. W. P. O’Meara, W.E. Collins, F.E. McKenzie, Parasite prevalence: A static measure of dynamic infections. Am. J. Trop. Med. Hyg. 77, 246–249 (2007).

54. R. De Jong, B. G. J. Knols, Selection of biting sites on man by two malaria mosquito species. Experientia 51, 80–84 (1995).

55. T. Dekker, et al., Selection of biting sites on a human host by *Anopheles gambiae s.s.*, *An. arabiensis* and *An. quadriannulatus*. Entomol. Exp. Appl. 87, 295–300 (1998).

56. L. Braack, et al., Biting behaviour of African malaria vectors: 1. where do the main vector species bite on the human body? Parasit. Vectors 8, 76 (2015).

57. A. Bompard, et al., High *Plasmodium* infection intensity in naturally infected malaria vectors in Africa. bioRxiv, 780064 (2019).

58. G. Valkiunas, Avian Malaria Parasites and other Haemosporidia (CRC Press, 2004).

59. J. Vézilier, A. Nicot, S. Gandon, A. Rivero, Insecticide resistance and malaria transmission: infection rate and oocyst burden in *Culex pipiens* mosquitoes infected with *Plasmodium relictum*. Malar. J. 9, 379 (2010).

60. R. Pigeault, et al., Avian malaria: a new lease of life for an old experimental model to study the evolutionary ecology of *Plasmodium*. Phil Trans R Soc B 370, 20140300 (2015).

61. M.J. Crawley, The R Book (John Wiley & Sons, 2012).

62. B.M. Bolker, Ecological Models and Data in R (Princeton University Press, 2008).

